# Evolutionary divergence of induced versus constitutive antiviral gene expression between primates and rodents

**DOI:** 10.1101/2024.05.26.595927

**Authors:** Lilach Schneor, Tzachi Hagai

**Affiliations:** The Shmunis School of Biomedicine and Cancer Research George S Wise Faculty of Life Sciences, Tel Aviv University, Tel Aviv 69978, Israel

## Abstract

**Background:** Hundreds of genes are upregulated in response to viral infection. These genes’ sequences often diverge across mammals, to counteract rapid virus evolution. However, the transcriptional divergence of these genes, their relative levels before and after infection in different host species, remains poorly understood.

**Results:** We studied this divergence by comparing gene expression before and after viral stimulation in cells from primates and rodents. We developed a method to identify orthologs upregulated in one species that are unchanged in response to stimulus in another species. Using human and mouse data, we detected 578 transcriptionally divergent orthologous genes. While most divergent genes do not belong to the same cellular process, several pathways and protein complexes are enriched in this set, suggesting that divergence in immune responses between closely related mammals is limited to specific modules rather than involving entire pathways. Transcriptional divergence between human and mouse orthologs was also observed when ortholog expression from different primates and rodents were compared, when responses were studied in several cell types, and was recapitulated at the chromatin level, using histone mark patterns that denote active promoter regions.

**Conclusions:** In summary, we found genes whose orthologs diverge between primates and rodents in response to immune stimulation. Some of these genes are constitutively expressed in one species even before infection, potentially facilitating rapid antiviral activity, and suggesting clade-specific adaptation to confer greater resistance against viruses. Further comparative studies on diverse infections can point to additional species-specific responses and how they enable different species to overcome infection.

## Introduction

The antiviral response is a cell-intrinsic innate immune program where hundreds of genes are rapidly upregulated in response to pathogen infection^1,2^. During the first wave of response, following sensing of pathogen-associated molecular patterns (PAMPs), such as dsRNA, numerous cytokines and chemokines are secreted from infected cells, to alert surrounding cells and immune cells of the infection^1^. Interferon (IFN) is often one of the cytokines secreted and its binding to IFN receptors on nearby cells leads to a second signaling cascade that results in the expression of hundreds of IFN-stimulated genes (ISGs) and to an antiviral state in both infected and uninfected cells^3–5^. Because many of the upregulated genes in both waves of response function directly or indirectly to inhibit infection, they are engaged in an evolutionary arms race with numerous viral proteins, leading to rapid evolutionary changes in the coding sequences of many antiviral genes that can alter their activity and the outcome of infection^6–9^. Unlike coding sequence evolution that has been characterized and mechanistically studied in numerous antiviral genes, the transcriptional evolution of antiviral genes and its potential role in host adaptation against viruses is less well-understood. Previous work that compared transcriptional responses to various PAMPs across species showed that genes that their level of response significantly changes between species have specific promoter architecture^10–12^. Several focused comparisons of this transcriptional response within clades, pointed to lineage-specific regulatory characteristics. For example, comparisons of the antiviral response of cells between bat two species^12^ and in lung infection across several mouse strains^13^ showed that genes upregulated only in specific species tend to be associated with either disease resistance or disease tolerance, reflecting different adaptation strategies to pathogens. Another study that compared antiviral and antibacterial responses in great apes and Old world monkeys showed that apes tend to upregulate a broader array of defenses in comparison to Old world monkeys^14^.

Here, we focus on a particular type of transcriptional divergence of antiviral genes between species, where orthologous antiviral genes are induced following immune stimulation in one species but remain transcriptionally unchanged in another species. In these cases, there are two general scenarios: (a) in one species the gene is upregulated in expression from a lower to a higher level following immune stimulation, while in the other species the orthologous gene remains constitutively low in expression, and (b) in one species the gene is upregulated, while in the other species the orthologous gene remains constitutively high before and after stimulation (see **Figure 1** for examples where the induced and constitutively expressed genes are human and mouse orthologs, respectively). The two scenarios can be related to two physiologically different states, depending on the level of expression of the orthologous genes that its expression remains unchanged: In the first scenario, the gene is not expressed or is lowly expressed in a given cell type and species before and after stimulation, and thus may not be part of the innate immune defense against pathogens in this species. In the second scenario, the gene is expressed in high levels before and after immune stimuli in one of the species, suggesting that this gene acts as a sentinel in this species, with its constitutively high expression level allowing for a rapid inhibition of viruses. The second scenario may thus represent an evolutionary adaptation where a gene evolved to be highly expressed in unstimulated cells and tissues to provide greater protection against an attack. This scenario was observed in *Pteropus Alecto,* the black flying fox, that its uninfected cells constitutively express IFNs or IFN regulatory factors, which is thought to confer a stronger protection against various viruses^15,16^. A similar “evolutionary switch” in gene transcriptional behavior was observed in the antiviral gene RNase L, that is strongly upregulated in response to IFN in black flying fox cells but is not upregulated in human cells^17^. Few examples of such transcriptional divergence in immune responses were also reported between human and mouse orthologs^10,11^, but a global analysis of this pattern of divergence is still lacking.

**Figure 1:**
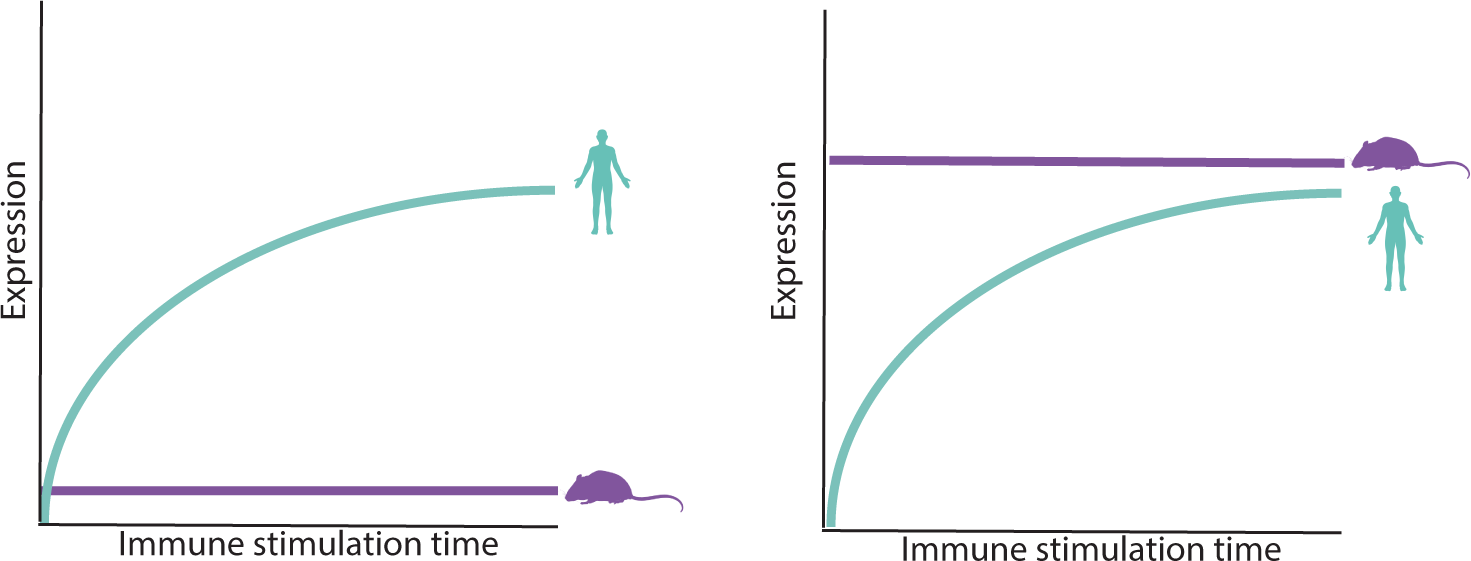
Schematics of the two scenarios of transcriptional divergence between species. In each panel we show the expression level over time of human and mouse orthologs, representing transcriptionally divergent genes we study in this work. In each of these panels, the human orthologous genes is induced following immune stimulation (such as following dsRNA or IFN treatment), while the mouse orthologous gene remains transcriptionally unchanged. The difference between the two scenarios is in the expression level of the mouse gene: In the left panel, the mouse gene level remains low or the gene is not expressed before and after stimulation. In the right panel, the mouse ortholog is already expressed in high levels before stimulation, and remains in this level following stimulation. The two panels represent functionally different scenarios: In the left panel, the mouse ortholog is not expressed and is not part of the immune defense program, unlike its human ortholog. In the right panel, the mouse ortholog is constitutively highly expressed, such that it can immediately function against invading viruses without the need to be induced, unlike the human ortholog that its levels are increased only after infection.

To comprehensively identify cases in which antiviral genes transcriptionally diverge and, specifically, switch between constitutive and induced expression across clades, we chose to use RNA-seq data from comparative cross-species *in vitro* systems, where homologous cells from different species belonging to several clades are stimulated and trigger an antiviral gene expression program: we focused on primate and rodent dermal fibroblasts stimulated with polyinosinic:polycytidylic acid (poly(I:C)), a synthetic dsRNA recognized by intracellular sensors, resulting in a strong upregulation of the antiviral response^18^. The upregulated genes include, among others, a diverse array of cytokines and chemokines, restriction factors that act to suppress viral replication, apoptotic factors and numerous genes related to the regulation of this response. Our analysis focuses on these genes’ relative expression before and after stimulation across a set of primate and rodent cells. We used two different, previously published, dsRNA-stimulated cross-species dermal fibroblast systems (**Figure 2A**): (1) A dataset that includes fibroblasts of two primates (human and rhesus macaque) and two rodents (mouse and rat) stimulated with dsRNA for 4 hours^11^, hereafter termed the **4-species stimulation system**. The data includes two additional datasets useful for our current analysis: analogous ChIP-seq data of H3K27ac histone marks, that denote active chromatin regions^20,21^, in both control and dsRNA stimulation across cells from the four species, and IFNB stimulation of cells from the same four species. (2) A second dataset of cells from nine primates (five great apes – human, chimpanzee, bonobo, gorilla, orangutan; three Old World monkeys – rhesus and pig-tailed macaques and baboon; and one New World monkey - squirrel monkey) and one rodent – mouse, unstimulated or dsRNA-stimulated for 24 hours^19^, hereafter termed the **10-species stimulation system**. As we demonstrate in the following analysis, in both 4- and 10-species stimulation systems the whole transcriptome and the transcriptional response to dsRNA tend to be more similar between closely related species, suggesting that regulatory changes gradually accumulate over evolutionary time, as observed in other expression programs^20–23^. This allows us to identify specific genes whose primate and rodent orthologs display the distinctive transcriptional divergence we focus on in this work: induced expression following dsRNA-stimulation in one clade and constitutive expression in the second clade. We then study the characteristics of these genes, including their cellular functions and coding sequence evolution, their expression in different human and mouse tissues and cells, and their transcriptional patterns in different species of primates and rodents.

**Figure 2:**
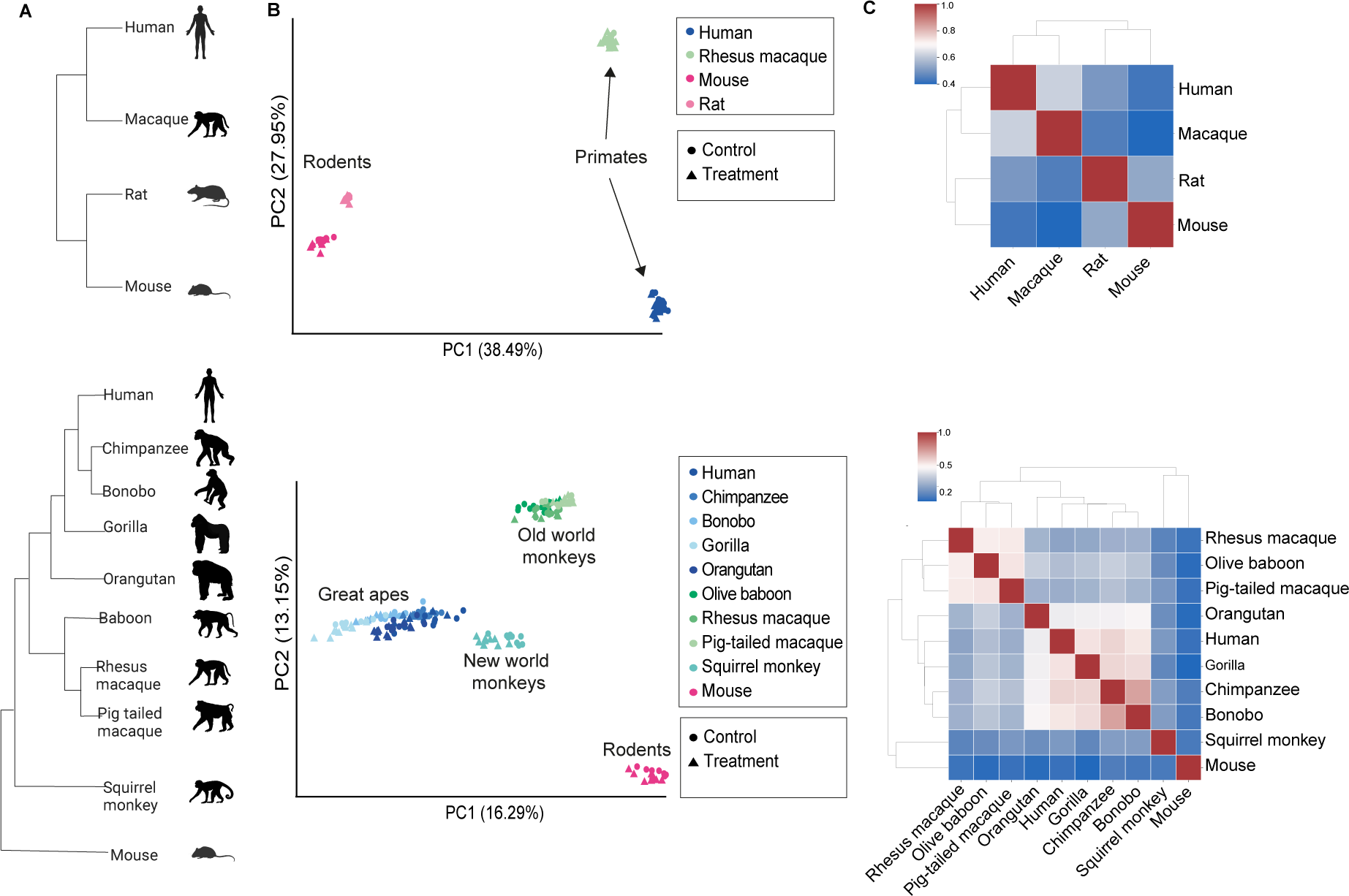
A cross-species *in vitro* immune stimulation analysis shows an overall similarity of transcriptional response across primate and rodent cells with accumaltive diveregnce. (**A**) Phylogenies of the two cross-species fibroblast dsRNA-stimulation systems used in this analysis. **Top**: 4-species stimulation system: Dermal fibroblasts from two primates and two rodents were profiled with and without dsRNA stimulation using RNA-seq, along with ChIP-seq of histone marks and IFNB stimulation profiling^11^. **Bottom**: 10-species stimulation system: Dermal fibroblasts from nine primates and mouse were profiled with and without dsRNA stimulation using RNA-seq^19^. (**B**) Scatter plot of principal component analysis (PCA) of expression levels (count values from Salmon mapping^48^). The proportion of variance explained by the principal components is indicated in parentheses. (**C**) Hierarchical clustering of pairwise Spearman’s rank correlations between fold change in response to dsRNA-stimulation values in 1-to-1 orthologous genes that are DE in at least one of the species (2,771 and 7,711 genes, top and bottom, respectively).

## Results

### Global cross-species analysis of gene expression and transcriptional response to dsRNA

To investigate the global patterns of gene expression similarity across species, we performed a principal-component analysis (PCA) based on the expression of one-to-one ortholog genes in RNA-seq libraries from the 4-species and 10-species datasets (12,811 and 11,350 orthologs, respectively). In each case we used all samples from this system (all individuals, samples from both stimulated and unstimulated conditions and including all species, **Figure 2B**). In both systems, the PCA plots show a separation by species rather than by condition. In each plot, we observed a separation that largely follows the phylogenetic relationship between the species: a separation between primates and rodents in both studies, and a separation between the major clades of primates in the 10-species dataset.

Next, we performed differential expression (DE) analysis between control and dsRNA stimulation samples within each species in each of the two studies. To study the overall similarity in response to dsRNA, we used the set of DE genes, defined as genes with an FDR-corrected P-value<0.01 in the DE analysis, in at least one of the species - 2,771 and 7,711 genes in the 4-species and the 10-species dataset, respectively. We compared the Spearman’s rank correlation coefficients of the fold change (**FC**) in response to dsRNA between species: For each pair of species, we computed the correlation between the FC values across the orthologous genes from the DE gene set, using the same set of genes for all pairwise comparisons. We then performed hierarchical clustering using the correlation values across all pairs of species (**Figure 2C**). We observed that species cluster in a manner that largely recapitulates their phylogenetic relationship. For example, in the 10-species dataset, the FC correlation values of the Old World monkeys first cluster with each other, similarly to the FC values of the great apes. These two clusters form one mega-cluster of all Catarrhini, leaving as outgroups the New World monkey and the rodent, squirrel monkey and mouse, as expected from their phylogeny.

Taken together, the data from both analyses, suggests that the overall gene expression (tested with PCA of orthologous gene expression) as well as the transcriptional response to dsRNA (tested with hierarchical clustering of FC values) are largely conserved between the set of studied species, and that they diverged in a similar manner to their known phylogeny, in agreement with previous analyses of transcriptional divergence data^19,21–23^. This overall similarity with accumulative divergence over evolutionary time, will allow us to focus on genes with particular patterns of divergence in dsRNA-response between the primate and rodent clades. Developing an approach to find these genes and their characterization will be the subject of the following sections.

### Identifying primate-rodent divergence between constitutive and induced antiviral gene expression

We next sought to identify genes whose orthologs’ expression differ between primates and rodents, such that in one clade the gene is induced following dsRNA stimulation, while in the other clade its expression remains largely unchanged. For this, we first focused on 1-to-1 orthologs between human and mouse from the 4-species dataset (16,534 genes). Using one-to-one ortholog gene expression data in dsRNA stimulation and control of human and mouse we devised the following approach to identify such genes (see **Methods** for details):

1. First, we performed a DE analysis between species. Unlike “regular” DE analyses, as in the previous section, where the DE is done between conditions within the same species, here we performed it by comparing gene expression between the human and mouse orthologs within the same condition: i.e., between human and mouse orthologs in stimulated conditions and, separately, between human and mouse orthologs in unstimulated control **(Figure 3A)**. This procedure yielded genes that differ in expression between human and mouse, in either dsRNA-stimulation or in unstimulated control (basal state). DE genes are here defined as those with an FDR-corrected P-value<0.001 and with a |Fold Change|>1 between the human and mouse orthologs in the tested condition **(Figure 3B)**.
2. Next, we excluded genes that differ between human and mouse in both dsRNA-stimulation and in control. In this manner, we filtered genes that are always higher in one species with respect to the other species, regardless of the antiviral stimulation (e.g., genes that are higher in human versus mouse cells, both in control and in dsRNA stimulation). This was achieved by keeping only genes that were found to be DE between human and mouse ortholog in one condition (as defined in Stage 1), but that their expression is not significantly different between human and mouse in the other condition (**Figure 3C**). We thus removed in this stage the set of genes that are less relevant to our study, such as those housekeeping genes that significantly differ between human and mouse, regardless of viral infection.
3. Finally, from the resulting filtered set, we focused only on genes that are significantly upregulated in response to stimulation in either human or mouse. This was done based on DE analysis between stimulation and control, within the same species (using the analysis in the previous section). These DE genes are defined as those with an FDR-corrected P-value<0.01 and that their FC>0). In this manner, we ensure that the final set of genes includes genes that viral stimulus significantly affects their expression in at least one species (**Figure 3C**).

**Figure 3:**
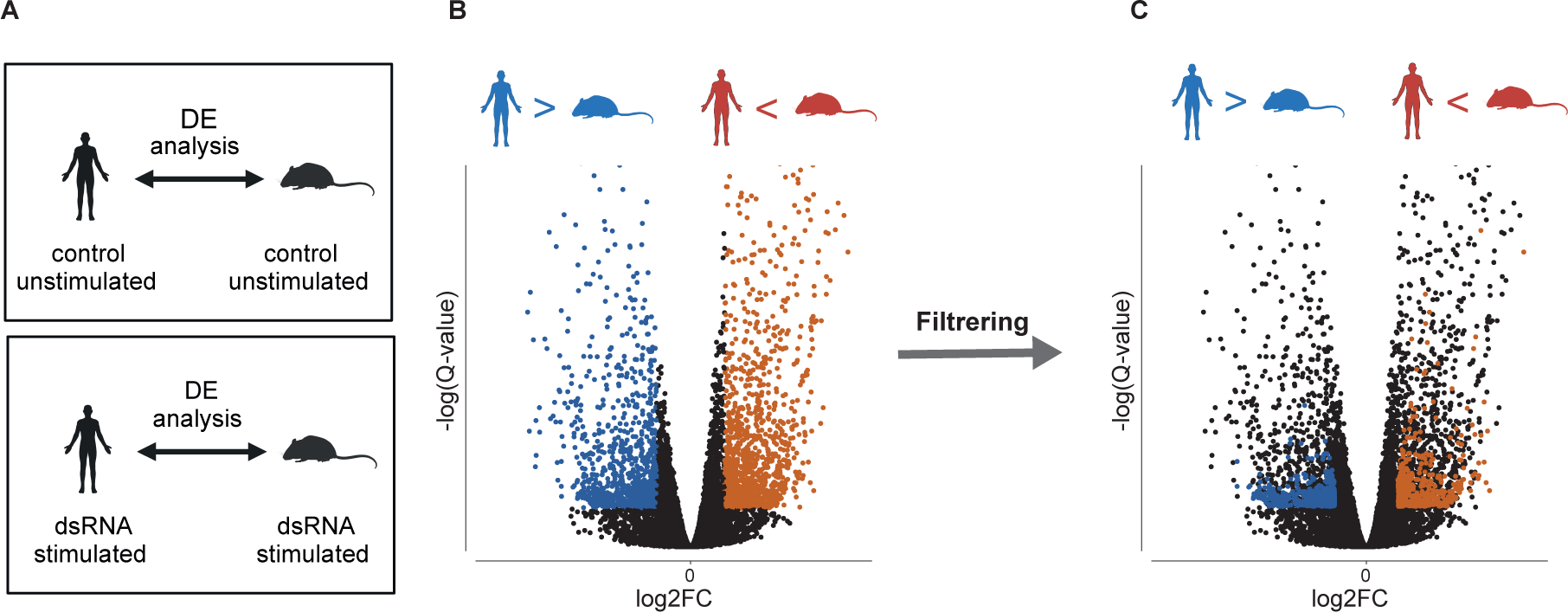
A method to identify divergent genes induced in one species and constitutive in the other. (**A**) Two separate DE analyses between human and mouse samples were performed in stage 1: One in the control-unstimulated conditions (top) and another in the dsRNA-stimulated conditions (bottom). (**B**) A representative volcano plot of DE analysis between ortholog gene expression of human and mouse in samples taken in the same condition, as described in stage 1. Each dot represents DE values (Fold Change and Q-value) of a one-to-one orthologous gene between human and mouse. Colored dots represent genes that differ in expression between human and mouse (FDR-corrected P-value<0.001 and with |Fold Change|>1), either in the control-unstimulated condition or in the dsRNA-stimulated condition. Orange dots are genes more highly expressed in mouse than in human, and blue are genes expressed more highly in human than in mouse in this specific condition. (**C**) We employed a filtering procedure to identify genes uniquely upregulated only in one species from the blue and orange sets: First, we filtered genes that differ between human and mouse in both dsRNA-stimulation and control-unstimulated conditions (these genes are genes that their divergence is unrelated to the antiviral response since they are higher in one species regardless of immune stimulation). Next, we excluded genes that are not significantly upregulated in response to dsRNA stimulation in either human or mouse (this ensures removal of genes that are insignificantly upregulated in response to dsRNA treatment in at least one species).

### Four groups of human-mouse transcriptionally divergent antiviral genes

The above-mentioned procedure yielded four groups of genes (**Figure 4**): Two sets of genes that differ between human and mouse in the stimulated conditions, but not in the unstimulated conditions, and two sets that differ between human and mouse only in unstimulated controls. Interestingly, each of the two DE analyses produced a set of genes that is only induced in human and a set of genes that is only induced in mouse, such that we have two sets with orthologs induced only in human and two with orthologs only induced in mouse. The differences between the two DE analyses lie in the constitutive expression of the orthologous gene: in one DE analysis, the constitutively expressed ortholog has high levels of expression relatively to its induced ortholog, whereas in the other DE analysis the constitutively expressed ortholog has low levels of expression (see **Figure 1** and **Figure 4, bottom**). There is an important physiological difference between constitutively high or constitutively low expression of an antiviral gene: If an antiviral gene is expressed highly before infection, it can confer an increased degree of protection against future pathogen infection, without the requirement to be induced. In contrast, constitutively low expression of an antiviral gene in a given species, both before and after stimulation, may suggest that in this species and tissue, the host defense has evolved to resist pathogens without this particular antiviral gene (unlike in the other species, where this gene is induced and may play a role in that defense).

**Figure 4:**
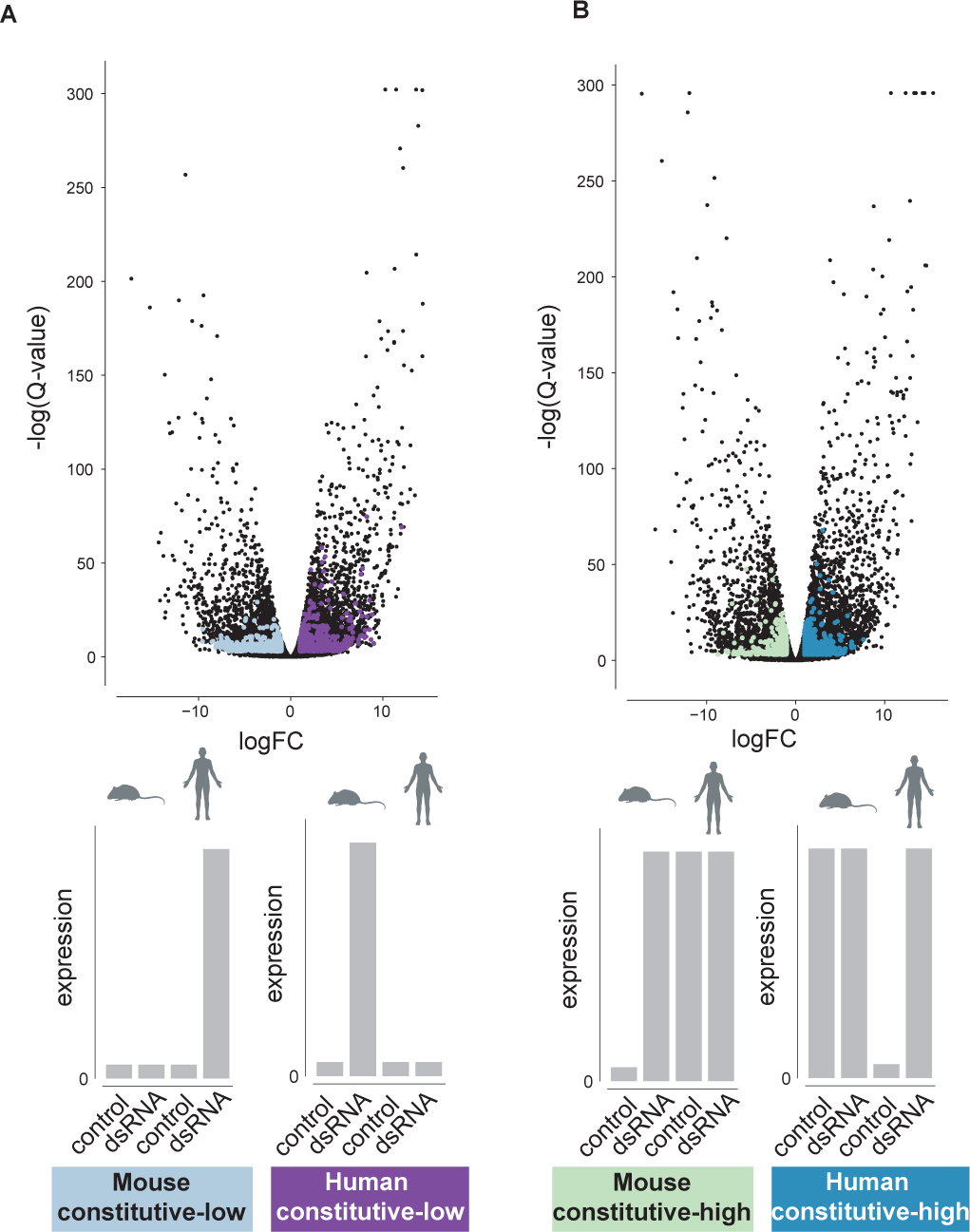
Four groups of human-mouse transcriptionally divergent genes and their patterns of expression. (**A**) A volcano plot of DE analysis between human and mouse in dsRNA-stimulated conditions. Colored dots show gene sets after the filtering procedure described in Figure 3. The right group (in purple) includes 224 genes induced in stimulation in mouse, but remain constitutively low in human (**‘human constitutive-low’**). The left group (light blue) includes 166 genes induced in stimulation in human, but remain constitutively low in mouse (**‘mouse constitutive-low’**). **Bottom**: Gray bars illustrate each group’s pattern of expression in human and mouse orthologs in both stimulated and unstimulated conditions. (**B**) The same as in A but comparing human and mouse in the control-unstimulated condition. The right group (turquoise) includes 104 genes induced in stimulation in human, but remain constitutively high in mouse (‘**mouse constitutive-high**’). The left group (light turquoise) includes 84 genes induced in stimulation in mouse, but remain constitutively high in human (**human constitutive-high**). **Bottom**: Gray bar plots illustrate each group’s pattern of expression in human and mouse orthologs in both conditions.

The human-mouse DE analysis in stimulated conditions resulted in two groups of genes: (1) a group of 166 genes induced in stimulation in human, but in mouse remain constitutively low (we term this group ‘**mouse constitutive-low**’), and (2) a group of 224 genes induced in stimulation in mouse, but remain constitutively low in human (‘**human constitutive-low**’) **(Figure 4A).** Similarly, the human-mouse DE analysis in unstimulated control resulted in two additional groups: (3) a group of 104 genes induced in human, whose expression levels in mouse are constitutively high (we term this group ‘**mouse constitutive-high**’), and (4) a group of 84 genes induced in mouse, that remain constitutively high in human before and after stimulation (‘**human constitutive-high**’) **(Figure 4B**). Detailed gene lists appear in **Supporting Table 1.**

Overall, these four sets of genes include 578 genes, out of 1,473 genes that are DE in either human, mouse or both. Thus, 39.2% of the DE genes in this cell system are induced only in one species and remain constitutively high or low in the other species.

### Expression levels of antiviral genes across human and mouse individual samples

Our procedure allows for identification of genes that are either up- or downregulated in one species in response to stimulation, while their expression in the other species is not significantly different between stimulation and control. To visualize the expression of these genes across species and conditions, we plotted heatmaps of their normalized expression (log10(TPM) - transcript per million) across all mouse and human individuals in both dsRNA stimulation and control conditions (**Figure 5**). Using hierarchical clustering, we observed that all individual from both species cluster together based on the observed pattern of expression rather than by species: For example, in “human constitutive-low” genes, we observed that the human samples, from both stimulated and unstimulated conditions, cluster with the mouse unstimulated samples, while the mouse stimulated samples are separated **(Figure 5A)**. This is consistent with our characterization of this set as genes that are constitutively low in mouse while being induced in humans, based on the cross-species DE approach described above. Thus, the unstimulated level of expression in human is similar to the level of expression of their mouse orthologs in both stimulated and unstimulated conditions.

**Figure 5:**
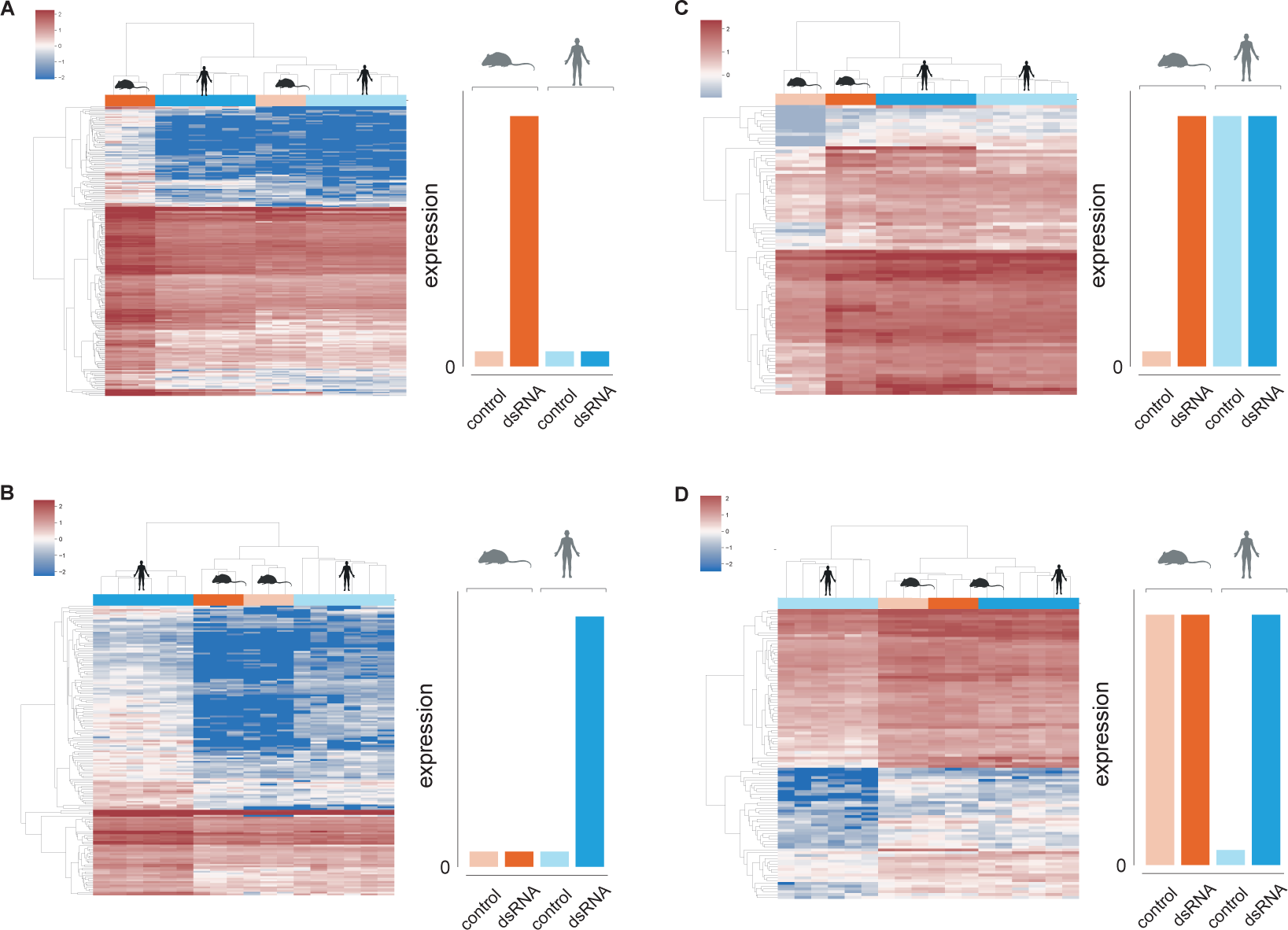
Hierarchically clustered heatmaps based on gene expression levels. Hierarchical clustering of gene expression data from all human and mouse individuals in the 4-species system in both stimulated and unstimulated conditions. In each panel, one the four previously defined divergent genes groups is shown, with expression levels in the human and mouse orthologs: (**A**) ‘human constitutive-low’, (**B**) ‘mouse constitutive-low’, (**C**) ‘human constitutive-high’ and (**D**) ‘mouse constitutive-high’. Columns are colored by species (blue and orange for human and mouse, respectively) and by conditions (dark and light for stimulated and unstimulated conditions, respectively) and represent one individual each. Gene levels are shown on a scale based on log(TPM). The clustering indicates that genes have similar levels in three ‘types’ of data, while the fourth is significantly different. For example, in A, both stimulated and unstimulated human samples cluster with the mouse unstimulated samples, while the stimulated mouse is the outgroup, with significantly higher levels of this gene set. This agrees with the patterns observed in Figure 4.

We also observed that the genes within each group are separated into two or three clusters. Each of these clusters represents a different level of relative expression. For example, in “human constitutive-low” genes, we observed two clusters of genes: In both clusters the level of expression in stimulated conditions in mouse samples is higher than the other three types of samples (unstimulated mouse samples, stimulated and unstimulated human samples). However, the higher cluster (in shades of blue to white) represents a set of genes that are lowly expressed and in stimulation of mouse cells increase in expression roughly by two folds. The lower cluster (in shades of light and dark red) has a similar pattern, but its gene levels are significantly higher (the basal expression levels are of logTPM of ∼1, while the induced mouse gene levels are of logTPM of ∼2). Thus, within each of the four sets of divergent human-mouse genes, we obtained several subsets of genes with different relative levels of expression in both unstimulated and stimulated conditions.

### Antiviral genes switching between induced and constitutive expression between primates and rodents are enriched in specific pathways and protein complexes

In the previous sections we identified groups of antiviral genes that switch between constitutive and induced expression when compared between human and mouse. We next asked whether these genes are enriched in particular pathways or functions. To this end, we employed GO term enrichment analysis, using the g:Profiler program^24^. As expected, the four sets are enriched in ‘general’ immune-related terms, such as those related to cytokine regulation and to response to virus. This is expected given their upregulated in response to dsRNA stimulation in at least one species.

In addition to these terms, in each of the four sets we also found enrichments that refer to more specific pathways or cellular functions, as well as genes that their protein products belong to the same protein complex, as defined by the CORUM database^25^, a collection of experimentally verified mammalian protein complexes. In the ‘human constitutive-high’ group, we found two genes - TAF1A, TAF1D, that are both members of the SL1 complex that is important for the assembly of the RNA polymerase I preinitiation complex^26,27^. Furthermore, the same set is enriched with genes associated with the TNF signaling pathway (see **Supporting Table 2**). In contrast, the ‘mouse constitutive-high’ group is enriched with genes associated with circulatory system development (including BMP7, KLF4, TNFAIP2 and FGF18). In the ‘mouse constitutive-low’ set, we found two members of the OAS pathway (a central pathway in viral RNA sensing^28^) - OAS2 and OAS3. In ‘human constitutive-low’ we found additional viral sensors and genes involved in upstream regulation of the innate immune response, the inflammatory response and IFN upregulation, including several NOD-like receptors and genes belonging to the inflammasome (NOD1, NLRC5, NLRP3)^29^, the NFkB complex (NFKB1, NFKBIE, IKBKE), as well as TRAF2, TBK1 and TANK that belong to the kinase maturation complex and are involved in NFkB signaling and cell survival^30,31^. Altogether, this suggests that there is no major antiviral gene pathway that all of its gene components shift in regulation between induced and constitutive gene expression across primates and rodents. However, such transcriptional changes occur within smaller sets of genes belonging to specific signaling or regulatory pathways and in particular modules and protein complexes.

### Coding sequence evolution and duplication rate of transcriptionally divergence antiviral genes

We next asked whether the identified antiviral genes that switch in regulation between constitutive and induced expression, also evolve rapidly in other evolutionary mechanisms including coding sequence evolution and fast rate of gene duplication and loss. For this, we compared the coding sequence evolution of the four sets of genes with the entire set of antiviral genes (defined as all genes that are DE in either mouse, human or both in response to dsRNA). When looking at sequence similarity between the human and mouse orthologs or at their dN/dS values (the ratio of non-synonymous to synonymous substitutions), we observed that the mouse constitutive-low group tend to evolve faster in sequence evolution, displaying lower sequence identity and greater dN/dS values (**Figure 6A-B**). This trend was also observed in the human constitutive-low group, but was not statistically significant. The human and mouse constitutive-high groups showed an opposite trend of higher sequence conservation than the group of all DE genes, that was stronger in the mouse group of genes.

**Figure 6:**
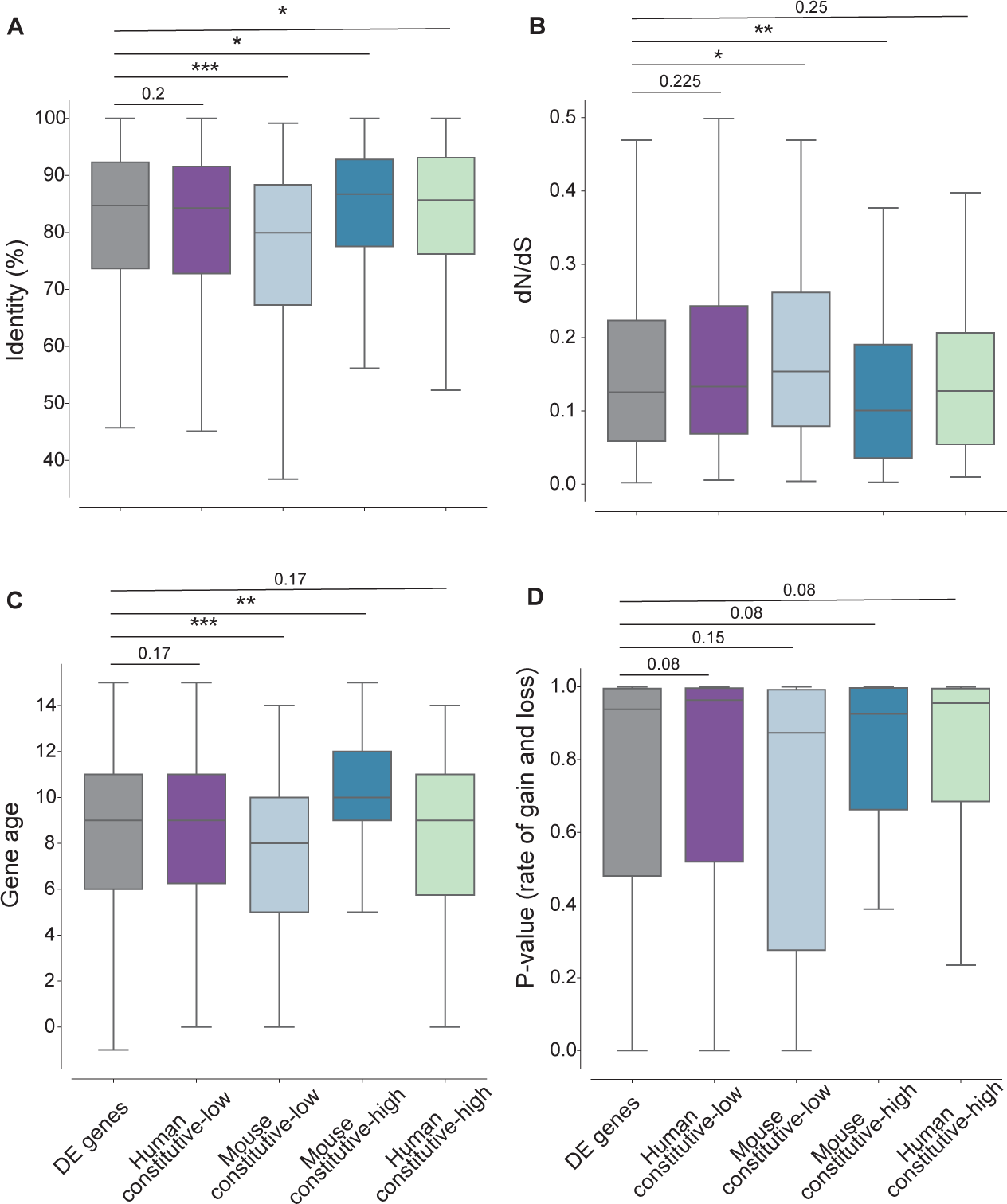
Evolutionary characteristics of genes that transcriptionally diverge between human-mouse in dsRNA response. Distributions of (**A**) percentage of sequence identity between human and mouse orthologs, (**B**) ratio of non-synonymous to synonymous substitutions (dN/dS), (**C**) gene evolutionary age (higher values denotes older age), and (**D**) rate of gene gain and loss across vertebrates (p-values for each gene’s rate are shown), for (from left to right): the set of DE genes (in grey) that includes all genes significantly upregulated in either human and/or mouse in response to dsRNA stimulation (FDR-corrected P-value<0.01 and FC>0), human constitutive-low, mouse constitutive-low, mouse-constitutive-high, human-constitutive high. Group colors, sizes and genes are as in Figure 4. FDR-corrected P-values are shown, one-sided Mann–Whitney tests were performed for each of the 4 groups against all DE genes.

When looking at the inferred evolutionary age of genes in the various sets, using age estimations from ProteinHistorian^32^, we observed that the mouse constitutive-low gene set was enriched with evolutionarily young genes while the group of mouse constitutive-high genes was enriched with evolutionarily ancient genes (**Figure 6C)**. The two groups of human constitutive high and low did not show significant differences in age distribution from the set of all DE genes. Finally, we did not observe significant differences in the rates of gene duplication and loss in the four sets versus the DE set (**Figure 6D**).

Thus, antiviral genes that exhibit a switch between constitutive and induced gene expression between human and mouse, show higher sequence divergence when the species-specific constitutive expression is low, and the opposite is true when the species-specific constitutive expression is high. These patterns are in agreement with previous findings on the relationship between gene expression and sequence conservation^33^.

### Human-mouse divergence in dsRNA response is recapitulated at the chromatin level

We next tested whether the expression patterns we observed at the transcriptional level are also observed at the chromatin level. For this, we used ChIP-seq data from the same system where the human and mouse dermal fibroblast response to dsRNA was profiled using RNA-seq. We focused on the histone mark H3K27ac, that is associated with higher activation of transcription^11,34,35^, and used H3K27ac ChIP–seq peaks in the vicinity of gene’s transcription start site (TSS) (peaks overlapping with a region 2,000bp upstream to 500bp downstream of the TSS), to define active promoter regions (see **Methods**). We compared the presence of these active histone marks in human and mouse cells in unstimulated and dsRNA-stimulated conditions in the four groups of genes we identified.

We reasoned that transcriptional activity across species and conditions should be reflected in patterns of active chromatin marks. For example, in the group of ‘mouse constitutive-high’ genes, the genes are highly transcribed in mouse in both stimulated and unstimulated conditions, but in human cells they are only highly transcribed during stimulation. In this case, histone marks should be present in both conditions in mouse, but only in the stimulated conditions in human cells, if the chromatin marks reflect the transcriptional behavior. To this end, we indeed observed this expected pattern in promoter regions of this gene set: an enrichment in the presence of the histone marks in mouse in both stimulated and unstimulated conditions and only following stimulation in human (FDR-corrected P-value = 0.0066, Fisher’s exact test, **Figure 7A**). This enrichment was observed in all four groups of genes and was statistically significant in three of the four (in ‘mouse constitutive-low’ the P-value is 0.44, Fisher’s exact test) (**Figure 7A**).

**Figure 7:**
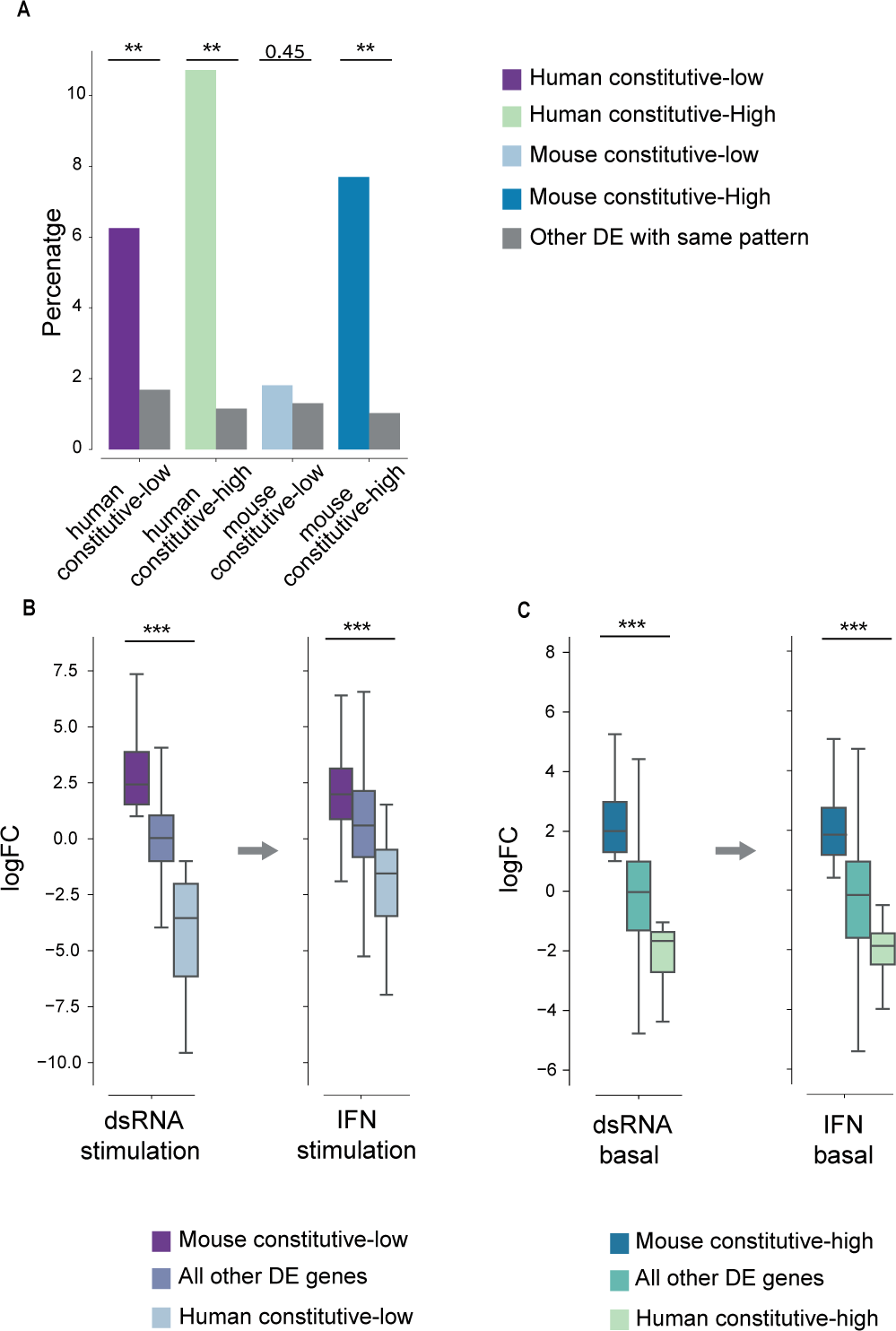
Human-mouse divergence in dsRNA response is recapitulated at the chromatin level, in IFN response and across different cell types. (**A**) Fraction of genes that their pattern of promoter activity agrees with gene expression, matching one of the 4 divergent groups (colored as in the legend) versus the fraction of genes among the rest of DE genes that display this pattern (in grey). For example, ∼6% of human constitutive-low genes show the expected pattern at the chromatin level, and ∼2% of all other DE genes show the same pattern (two left-most bars). FDR-corrected Fisher’s exact test P-values are shown. These tests show that the pattern of transcriptional response is recapitulated at the chromatin level in all four groups since the fraction of genes whose chromatin accessibility pattern agrees with the transcriptional pattern is higher than that observed in the set of all other DE genes (this is significant in three of four groups tested). (**B**) logFC distribution values based on DE analyses between human and mouse in dsRNA stimulation (before the arrow) and in IFN stimulation (after the arrow). In both cases the genes are partitioned into three boxplots based on three groups defined previously: “human constitutive-low”, “mouse constitutive-low” and “all other DE genes”. Thus, the same three groups of genes are shown before and after the arrow, but their FC values varies based on either dsRNA- or IFN-stimulation DE values. The analysis shows that the separation in logFC values in the dsRNA-stimulation between the three groups is also observed in the IFN-stimulation, suggesting similar transcriptional divergence between dsRNA response and IFN response. **(C)** The same as in B, but with basal conditions used in the DE analyses and showing: “mouse constitutive-high”, ‘human constitutive-high”, “all other DE genes”. In both B and C, FDR-corrected P-values are shown for one-sided Mann–Whitney tests, performed under the hypothesis that the FC distribution of the left group is higher than the right group.

We note that a fraction of the genes in these sets has ChIP-seq peaks in all four states or in none of them (human and mouse cells, stimulated and unstimulated conditions). This is also true for many of the DE genes in general, and either reflects smaller dynamics at the chromatin level than at the transcriptional level (where the chromatin remains accessible even without transcription), or the detection and quantification limits using ChIP-seq. In summary, the divergence observed between human and mouse in transcriptional response to dsRNA is also reflected to a large and significant extent at the chromatin level.

### Human-mouse divergence in dsRNA response is reflected in secondary response induced by interferon

Next, we compared human and mouse dermal fibroblast response to interferon (IFN), using stimulation data obtained in parallel to the dsRNA stimulation in the original study^11^. IFN is rapidly secreted following viral or bacterial infection, leading to a strong secondary wave of response against pathogens, including upregulation of hundreds of interferon stimulated genes (ISGs), some of which are also induced in response to dsRNA. Similarly to the analysis of the human-mouse dsRNA stimulation system (shown in **Figures 3-4**), we performed DE analysis to compare between human and mouse cells in unstimulated conditions and, separately, comparing IFN-stimulated human and mouse cells. We then tested whether genes previously detected to diverge between human and mouse in dsRNA, diverge in a similar manner in IFN response (e.g., genes that were defined as “mouse constitutive-low” in the dsRNA stimulation analysis would behave in a similar manner in unstimulated and IFN-stimulated cells). For this, we tested whether the resulting fold change in ortholog expression in the IFN system is similar to what was found between human and mouse in the dsRNA system. We observed that transcriptional trends in response to IFN in human and mouse largely and significantly match those found in the dsRNA response (**Fig 7B-C**, **Supporting Figure 1**).

For example, when splitting the genes in the IFN response to four classes, as done previously with the dsRNA stimulation data, we observed that genes originally defined as “human constitutive-low” based on the dsRNA system have the highest FC values in the IFN system while the “mouse constitutive-low” genes have the most negative FC values (**Figure 7B**). These trends suggest similarities in transcriptional divergence between the dsRNA and IFN systems that were also observed in the sets defined as “mouse constitutive-high” and “human constitutive-high” (**Figure 7C**). Additionally, when separately using the TPM values in human and mouse replicates, we observed that the expression patterns observed in the dsRNA stimulation in the four-divergent groups between human and mouse was largely reflected in the IFN stimulation (**Supporting Figure 1**).

From these results we conclude that transcriptional divergence between human and mouse in IFN stimulation largely follows the same patterns as those originally observed in dsRNA stimulation. This serves as a further validation for the divergence we originally found in these genes transcriptional responses dsRNA between human and mouse.

### Human-mouse divergence in dsRNA response is recapitulated in other primate and rodent species

Next, we asked whether the divergence observed between human and mouse genes in dsRNA-response represents an overall divergence in dsRNA transcriptional responses between the primate and the rodent clades. To test this, we first repeated our analysis with dsRNA stimulation using the rhesus macaque and rat data from the same 4-species system we took the human and mouse data from. As in the human-mouse dsRNA stimulation analysis, we contrasted the macaque and rat 1-to-1 orthologs in the same condition (where samples from both species are either stimulated with dsRNA, or unstimulated). We then tested whether the resulting fold change in ortholog expression (between macaque and rat) is similar to what we found between human and mouse: In the comparison of macaque-rat dsRNA stimulated samples, we again divided the genes based on their orthologous human-mouse gene diveregnce and comapred their FC in response to dsRNA in the macaque versus rat comparisons. We observed that in the comparison of FC between rat and macaque, the group of genes originally defined as ‘human constitutive-low’ has the highest FC values between rat and macaque while the group of ‘mouse constitutive-low’ genes has the most negative FC values (**Figure 8A Top**). We observed the same trend when contrasting rat and macaque samples in control conditions and using the gene sets originally defined as ‘mouse constitutive-high’ and ‘human constitutive-high’ (**Figure 8A Bottom**). This suggests that the differences in constitutive versus induced expression of antiviral genes originally found between human and mouse are consistent with the expression patterns of their orthologous genes in macaque and rat.

**Figure 8:**
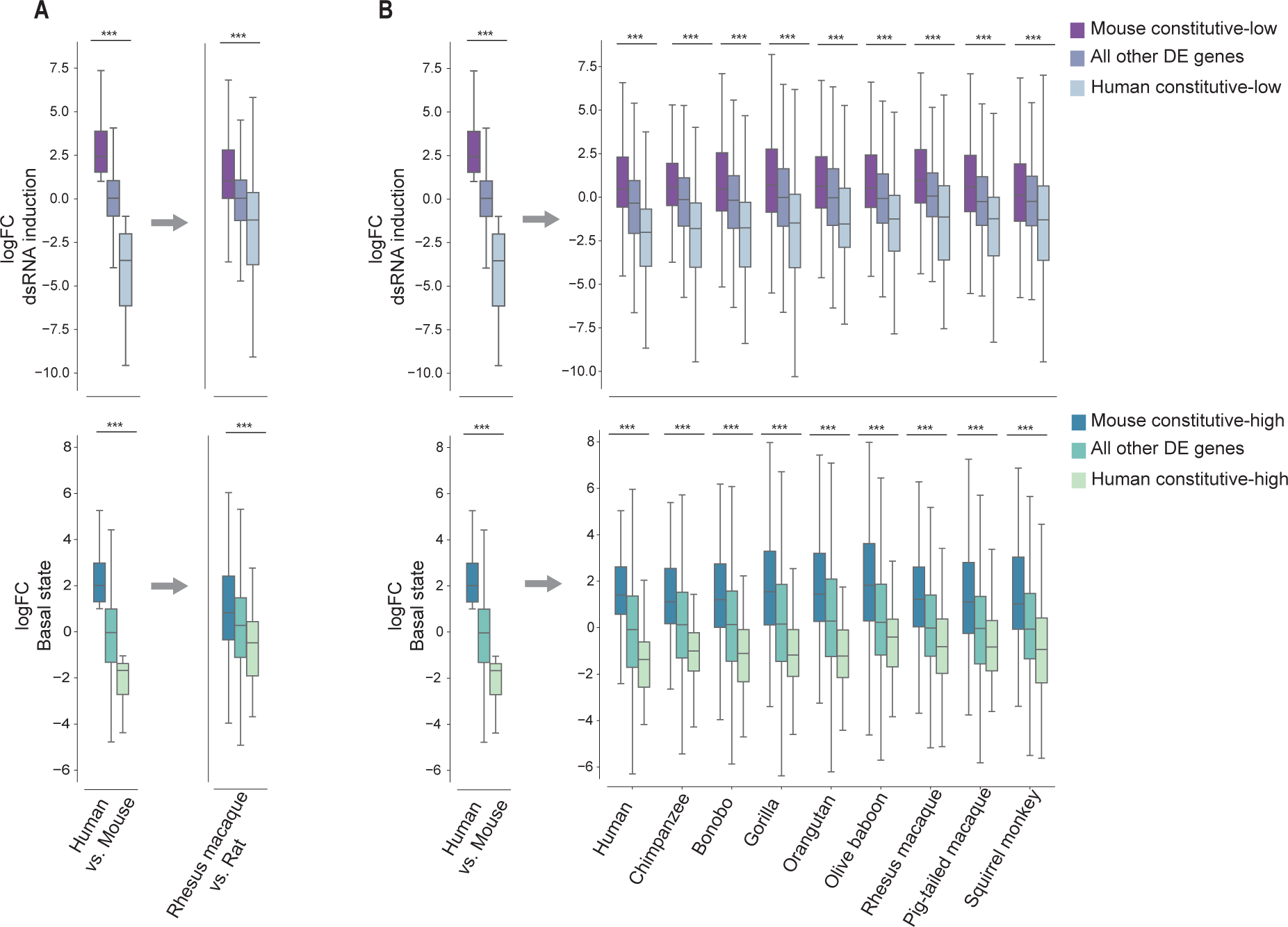
Divergence in dsRNA response across different primate and rodent species. (**A**) **Top**: logFC distribution of differential expression analysis in dsRNA-stimulation condition between human and mouse (left) and rhesus macaque and rat (right) from the 4-species system. Separation in both boxplots is according to the groups originally defined from human-mouse divergence: “mouse constitutive-low”, “human constitutive-low”, “All other DE genes” (as defined in Figures 3-4). The right panel, which is based on rhesus versus rat data shows that the orthologs of two species diverge in a similar manner to the human and mouse orthologs. Thus, both left and right panels use the same three groups of orthologous genes, but the FC values are based on either human-mouse (left) or macaque-rat (right) DE analysis. **Bottom**: The same as in Top, but with DE preformed between human and mouse in basal conditions and showing “human constitutive-high”, “mouse constitutive-high” and “all other DE genes”. (**B**) The same as in A, but comparing the human-mouse divergence in response to dsRNA from the 4-species system to each one of the 9 primates-mouse divergence in response to dsRNA from the 10-species system (the primate species used is shown in the bottom, and its data is always compared to mouse). In both (A) and (B) we observed that the patterns of relative FC between the compared groups recapitulate the patterns from the human-mouse dsRNA comparison. In all analyses, FDR-corrected P-values are shown for one-sided Mann– Whitney test, performed under the hypothesis that the FC distribution of the left group is higher than the right group in each of the analyses.

We further tested the transcriptional human-mouse divergence in the dataset that includes nine primates and mouse stimulation (the 10-species system). We repeated the same analysis as done with the rhesus and rat (**Figure 8A**) with the 10-species system (**Figure 8B**). In this system, we compared the mouse data to each of the 9 primates. In each one of the primate-mouse comparisons we again observed the expected trend, of higher FC values in the ‘human constitutive-low’ group in the dsRNA stimulation condition, and in the ‘mouse constitutive-high’ group in the control condition. This trend was also observed when looking at the lower FC values that were in the ‘mouse constitutive-low’ in the dsRNA condition and in the ‘human constitutive-high’ in the control condition **(Figure 8B).** This suggests that the transcriptional differences originally observed between human and mouse in response to dsRNA stimulation are conserved in other primates compared to rodents.

Finally, we asked whether the genes found to be diveregnt between primates and rodents, also change between different primate species based on the primate phylogeny. For this, we clustered the logFC values of the orthologous genes in each of the 10 species (nine primates and mouse), in the four groups of divergence (**Supporting Figure 2**). We did not find groups of genes that show strong change in behaviour across the primates (for example, that they are only highly uprgeulated in Great apes and not in other primates). However, we do see that the FC values in response to dsRNA stimulation for each of the four human-mouse divergent groups are clustered in a manner that largely recapitulates the evolutionary relationship between primates **(Supporting Figure 2)**. This suggests that while overall the induced versus constitutive expression behavior is consistent across primates, the levels of induction diverge to some extent between primates, and this follows the primate phylogeny.In other words, genes found to be induced following dsRNA stimulation in human but not in mouse are generally induced also in other primates, and the level of upregulation may be reduced as a function of the primate’s phylognetic distance from human.

### Human-mouse transcriptional divergence is consistent across different cell types

All previous analyses were performed on dermal fibroblasts that provide a unique system as they are a homogenous and comparative cell system across species, where data of dsRNA stimulation exists for numerous species. We next sought to expand the findings to other cell systems. While other cell systems are more limited in their data comprehensiveness, they can still offer comparisons for specific aspects in our study. We focused on two aspects of our findings: (1) the level of gene expression in non-induced cells and tissues, and (2) the innate immune induction following stimulation in various cells and tissues.

First, since for each species we have genes denoted as ‘constitutive-low’ or ‘constitutive-high’ based on basal expression in fibroblasts, we tested the basal expression levels of these two groups of genes across tissues, using expression data from a large set of mouse and human tissues (taken from the GTEx^36^ and the BodyMap^37^ datasets). We observed that genes defined as ‘constitutive-high’ based on the fibroblast data are significantly higher than ‘constitutive-low’ genes across tissues. This is true both for most human tissues (35 of 40 tested tissues, based on GTEx) (**Supporting Figure 3**) and for all mouse tissues (18 mouse tissues, based on BodyMap dataset) (**Supporting Figure 3**). This suggests that the basal expression of the identified genes in fibroblasts is largely consistent with relative basal expression levels of these genes across tissues.

Secondly, we asked whether transcriptional response patterns observed in the fibroblasts system, are also observed in other cell types. For this we utilized the Interferome database, that includes data from a wide range of IFN stimulation studies of various human and mouse cell types and tissues stimulated in various conditions and timepoints^38^. We note that such a comprehensive data does not exist for dsRNA stimulation. Using the human-mouse IFN analysis (described in **Figurer 7B**), which is based on dermal fibroblast IFN stimulation, we tested the generality of our results, in terms of divergence in response to IFN between human and mouse, in various cells and tissues appearing in the Interferome database. We took 27 human and 10 mouse stimulation experiments and tested for each gene in how many of these experiments it was significantly induced following Type-I IFN treatment. We divided the genes into the four transcriptionally divergent groups previously defined based on our human-mouse fibroblast IFN stimulation dataset. In all four transcriptionally divergent gene sets we observed: (1) a higher fraction of induction in one species than the other, and (2) the induction was higher in the species expected to show an induction based on the fibroblast system (**Supporting Figure 4**). For example, in the set of genes we originally defined as ‘mouse constitutive-high’ and ‘mouse constitutive-low’, we expected to see that more genes would be induced by IFN in human rather than mouse cells, if the patterns observed in fibroblasts are consistent across other cell types. This is is indeed the case, as observed in the higher fraction of IFN stimulation studies that detected these genes as IFN-induced. Additionally, in both ‘human constitutive-high’ and ‘human constitutive-low’ genes, we observed a higher fraction of induction in mouse cell systems, again suggesting a conserved pattern of IFN induction in a species-specific manner across cell types.

Thus, the divergence in induction following IFN stimulation between human and mouse fibroblasts is also observed in other human and mouse cell types. This suggests that species-specific induction is consistent across different cells and tissues, at least to some extent. We note however that the fraction of observed induction across cell types is lower than 30% (as can be observed by the median values of the distributions in **Supporting Figure 4**). Out of ∼16,000 1-to-1 orthologs assayed in these studies, 7,136 genes are upregulated in at least one study, while 103 are upregulated in at least half of the assays in both human and mouse systems. Thus, while species-specific induction is more conserved than expected, as reflected in the significant similarity to our fibroblast results across the four sets of genes defined in (P-values are always significant), there are still many genes that their induction following immune stimulation varies between different cell types and tissues. Moreover, our analysis suggests that there is a small core of ∼100 IFN-induced genes that are strongly upregulated across numerous tissues in both human and mouse, in agreement with previous results regarding conserved ISGs across mammals^4^.

## Discussion

In this work we aimed to identify cases in which antiviral genes switch between constitutive and induced expression across clades and species. The expression level of antiviral genes is often increased following immune stimulation or infection, to enable better protection against and restriction of invading pathogens^2,3,39^. However, most antiviral genes are also expressed in ‘basal conditions’, without infection or immune stimulation. Basal expression is thought to be important in providing an immediate protection, or readiness, against viruses. Both basal and induced levels of antiviral genes are under strict regulation, since excessive antiviral gene expression can lead to immune pathologies while reduced levels can compromise host defenses^40–44^. Evolutionary changes where antiviral genes are strongly induced in one species but are constitutively expressed in another species, may thus impact different host species abilities to inhibit various virus replication.

To study such “evolutionary switches” in antiviral gene expression, we focused on the comparison between primates and rodents – two biomedically important clades that have relatively large amount of comparative data both between and within the clades. We used two comparative *in vitro* systems of dermal fibroblasts including cells from several primates and rodents (4-species and 10-species systems) that were stimulated with dsRNA and include gene expression of both dsRNA-stimulated and control-unstimulated conditions. We first analyzed the whole-transcriptome and the overall response similarity between species, showing that the transcriptional divergence is overall similar to the phylogenetic relationship between these species, suggesting that most genes are conserved in expression and that divergence is accumulated over evolutionary time. This gradual divergence in the majority of gene expression across species, enabled us to focus on specific genes that significantly change in transcriptional response between primates and rodents.

Using a new approach to identify genes that differ in expression between primates and rodents, and first focusing on the comparison between human and mouse from the 4-species stimulation system, we generated four groups of human-mouse transcriptionally divergent genes. In two of those sets the genes are induced only in mouse, whereas in the other two, the genes are only induced in human. The difference between the two sets depends on whether the group of constitutive genes remains transcriptionally low or stays at relatively high expression levels in both control and dsRNA-stimulation conditions (see **Figure 1**). Our approach can be extended to studying other clades or other gene expression programs, and can be useful to detect such inter-species switches in transcriptional behavior in various cellular contexts.

We next characterized the functional characteristics of these four sets of genes. We found that each of the four sets is enriched with genes belonging to specific pathways or that their protein products are part of the same protein complex. However, our results did not point to a major pathway whose expression was completely altered between primates and rodents. This suggests that the divergence in innate immune response between these two closely related clades has occurred in specific regulatory changes of one or few genes, rather than involving a shift of an entire pathway. This may mirror findings of positive selection in antiviral genes, that showed that evolutionary arms races between host and their viruses are often associated with changes in specific residues of a few antiviral genes related to the restriction of the studied virus^45,46^.

Our evolutionary analysis of coding sequences of the transcriptionally divergent antiviral genes, showed that some of these groups tend to more conserved in sequence than the whole set of antiviral genes whereas others tend to be less conserved. These patterns may be related to the expression levels of these genes across tissues in basal conditions. In addition, they may reflect the different functions of the genes in these groups. This is in agreement with previous findings that showed that different classes of antiviral genes have different evolutionary rates and that only certain classes of antiviral genes are enriched with genes with signatures of positive selection^11,47^.

To test the generality of our results of the human-mouse dsRNA stimulation, we first compared the response to dsRNA and IFN stimulation. We found that the transcriptional divergence between human and mouse response to dsRNA was similar to the one in the IFN stimulation. This suggests a similarity in the divergence between the primary and secondary innate immune responses. The generality of our findings was also reflected to some extent when we compared the response to IFN of human and mouse fibroblasts to other human and mouse cell types, as observed in the analysis using the Interferome database. However, as expected, many genes vary in their induction between different cell types, reflecting cell-type specific response to infection. Looking at the conservation of response across different primates, we found that the majority of genes induced in human but not in mouse are also induced in other primates. We also found that the response within primates gradually diverge with evolutionary distance.

Taken together, our results point to a group of genes that diverge between primates and rodents in basal and induced expression in response to viral stimulation. The induction of these genes is largely recapitulated across the primate and rodent clade, and is shown to be in agreement with chromatin accessible regions in nearby promoter regions and, to some extent, in different cell types and tissues. While our work suggests that some of the genes that switch in transcriptional behavior in fibroblasts between primates and rodents also seem to behave similarly in other cell types, further comparative studies would be needed to fully establish this.

Together with previous studies focusing on identification of conserved immune-stimulated genes in mammals or primates^4,19^ and on different aspects of transcriptional divergence in immune response to infection^11–14^, our study contributes to understanding of the evolutionary landscape of the innate immune response and its regulation. The transcriptionally divergent genes found in this work can contribute to species-specific adaptation to viral infection and provide directions to understanding the mechanisms behind this adaptation.

## Methods

### Dataset assembly and availability

All data used in this work is publicly available, as detailed below. For comparing innate immune responses across species, we searched for comparative cell systems where similar cells grown in similar conditions were treated with the same immune stimulant to elicit an antiviral response across species. The two systems we used were both dermal fibroblasts, grown *in vitro* and stimulated with dsRNA (poly I:C) from primates and rodents: the 4-species system^11^ (**Arrayexpress accession - E-MTAB-5919**), and the 10-species system^19^ (**SRA accession: SRP120495**). The first system included 4 species - 2 primates (human and rhesus macaque) and 2 rodents (mouse and rat), stimulated with dsRNA for 4h, while the second included 9 primates (5 Great apes – human, chimpanzee, bonobo, gorilla, orangutan; 3 Old world monkeys – rhesus and pig-tailed macaques and baboon; and one New world monkey - squirrel monkey) and mouse, stimulated with dsRNA for 24h. Each of these systems included stimulation of cells from several individuals per species (only females in the 4-species system and both males and females in the 10-species system). Samples from each individual included treated (dsRNA) and mock treated control, profiled using RNA-seq.

In the case of the 4-species system, we also used IFNB stimulation and control for the same species and individuals, stimulated and processed in parallel to the dsRNA transfection. The 4-species system also included ChIP-seq data of histone marks, from which we used the H3K27ac data that marks active chromatin regions, to study active promoters (**ArrayExpress accession: E-MTAB-5918**).

We chose not to use several other cross-species fibroblast system studies^4,12^, since they lack more than a single primates and/or rodent species, required for our analysis.

### Read mapping and gene expression quantification

Reads were mapped and gene expression was quantified using Salmon (version 0.13.1)^48^ with the following commands:

For paired end libraries: “salmon quant -i {index_file_directory} -l A -p 8 --validateMappings - -gcBias --seqBias --numBootstraps 100 -g {transcript_to_gene_file} -1 {first_read} -2 {second_read}”. -o {output_directory}”

For single end libraries: “salmon quant -i {index_file_directory} -l A -p 8 --validateMappings -- gcBias --seqBias --numBootstraps 100 -g {transcript_to_gene_file} -r {read} -o {output_directory}”

Each sample was mapped to its respective species’ annotated transcriptome (coding genes only, downloaded from ENSEMBL version 99^49^: Human - GRCh38, Chimpanzee - Pan_tro_3.0, Bonobo - panpan1.1, Gorilla - gorGor4, Orangutan - PPYG2, Olive baboon - Panu_3.0, Rhesus macaque - MMUL_1, Pig-tailed macaque - Mnem_1.0, Bolivian squirrel monkey - SaiBol1.0, Mouse - GRCm38, Rat - Rnor_6.0). We removed annotated secondary haplotypes of human genes by removing genes with ‘CHR_HSCHR’.

### Quantifying differential gene expression in response to dsRNA and to IFNB

To quantify differential gene expression between treatment and control for each species and for each treatment separately, we used edgeR (version 3.32.1)^50^, using R version 4.0.3, using the rounded estimated counts from Salmon. Differential expression analysis was performed using the edgeR exact test, and P-values were adjusted for multiple testing by estimating the false discovery rate (FDR), similarly to our previous work^11,12^.

### Principal component analysis (PCA)

For PCA analysis we used the prcomp() function in R version 4.0.3, and as input the rounded estimated counts from Salmon. The analysis included only 1-to-1 orthologous genes in all of the species in the study (12,811 and 11,350 orthologs, for the 4-species and 10-species studies, respectively). Before performing the analysis, we transformed the data with log10 transformation on the counts values, and scaled it using the scale() function in R (center=True, scale=False).

### Hierarchical clustering of gene expression and fold change values

We used clustermap() function from seaborn package in Python version 3.9.7. Clustering parameters: method=’average’, metric=’euclidean’.

### Identification of genes dsRNA-upregulated in one species and constitutively expressed in another species

We performed the following procedure to identify genes that are upregulated in response to dsRNA in one species, but are constitutively expressed (do not significantly change in expression before and after treatment) in the other species.:

1. Differential expression analysis between the human-mouse orthologs <u>within the same</u> <u>condition</u> (dsRNA-stimulation conditions between human and mouse samples, and separately, the control-unstimulated conditions between human and mouse samples). DE analysis was performed using edgeR (version 3.32.1), with the rounded estimated counts obtained from Salmon. This was done only for genes that had a significant level of expression in at least 3 replicates (TPM>0, transcripts per million). Differential expression analysis was performed using the edgeR exact test, and P-values were adjusted for multiple testing by estimating the false discovery rate (FDR).
2. Identification of differentially expressed genes between human and mouse in either dsRNA-stimulation, control or both conditions: These DE genes are defined as those with an FDR-corrected P-value<0.001 and with a |Fold Change| > 1 between the human and mouse orthologs in the tested condition (thus, we obtained genes that are higher in expression in this condition in either human or mouse, depending on the sign of the FC).
3. For each of the groups of DE genes identified (with FC>1 or FC<-1 and FDR-corrected P-value<0.001, and in each of the two conditions), we removed genes that displayed the same behavior in the other condition. For example, genes that were significantly higher in mouse than in human cells (FC>1 and FDR-corrected P-value<0.001) in both the dsRNA-stimulation as well as in control-unstimulated conditions were removed (since they likely represent a set of genes that is higher in mouse versus human regardless of the innate immune response).
4. We further removed any gene that is not DE in response to treatment in at least one species. That is, if a gene was not found to have a q-value<0.01 and FC>0, in either human and/or mouse in response to dsRNA, it was excluded from the analysis (in this case, we used DE values obtained from a “classical DE analysis” where samples from the same species from two different conditions were compared to find DE genes in response to dsRNA-treatment in either human or mouse.

This resulted in 4 groups of species-specific responding genes (detailed lists appear in **Supporting Table 1)**:

1. ‘**Mouse constitutive-low**’: Generated from the group of genes that were significantly higher in human than in mouse in the dsRNA-stimulation conditions (FC<-1 and FDR-corrected P-value<0.001 in the DE analysis in dsRNA-stimulation conditions) (Stage 2). From this group we excluded genes that were also significantly higher in human than in mouse in the control-unstimulated conditions (FC<-1 and FDR-corrected P-value<0.001 in the DE analysis in the control-unstimulated condition) (Stage 3). We then removed genes that were not DE in response to treatment in at least one species (Stage 4). This resulted in 166 genes that are induced in dsRNA-stimulation only in human but in mouse remain constitutively low in both unstimulated and stimulated conditions.
2. ‘**Human constitutive-low**’-Generated from the group of genes that were significantly higher in mouse than in human in the dsRNA-stimulation conditions (FC>1 and FDR-corrected P-value<0.001 in the DE analysis in the dsRNA stimulation conditions) (Stage 2). From this group we excluded genes that were also significantly higher in mouse than in human in the control-unstimulated conditions (FC>1 and FDR-corrected P-value<0.001 in the DE analysis in the control-unstimulated condition) (Stage 3). We then removed genes that were not DE in response to treatment in at least one species (Stage 4). This resulted in 224 genes that are induced in stimulation in mouse but in human remain constitutively low in both unstimulated and stimulated conditions.
3. ‘**Mouse constitutive-high**’-Generated from the group of genes that were significantly higher in mouse than in human in the control-unstimulated conditions (FC>1 and FDR-corrected P-value<0.001 in the DE analysis in the control-unstimulated conditions) (Stage 2). From this group we excluded genes that were also significantly higher in mouse than in human in the dsRNA-unstimulated conditions (FC>1 and FDR-corrected P-value<0.001 in the DE analysis in the dsRNA-unstimulated conditions) (Stage 3). We then removed genes that were not DE in response to treatment in at least one species (Stage 4). This resulted in 104 genes that are induced in stimulation in human but in mouse remains constitutively high in both unstimulated and stimulated conditions.
4. ‘**Human constitutive-high**’-Generated from the group of genes that were significantly higher in human than in mouse in the control-unstimulated conditions (FC<-1 and FDR-corrected P-value<0.001 in the DE analysis in the control-unstimulated condition) (Stage 2). From that group we excluded genes that were also significantly higher in human than in mouse in the dsRNA-unstimulated conditions (FC<-1 and FDR-corrected P-value<0.001 in the DE analysis in the dsRNA-unstimulated conditions) (Stage 3). We then removed genes that were not DE in response to treatment in at least one species (Stage 4). This resulted in 84 genes that are induced in stimulation in mouse but in human remain constitutively high in both unstimulated and stimulated conditions.

### Analysis of basal gene expression across tissues in human and mouse

To characterize relevant gene expression across human and mouse tissues, we extracted the data from Genotype-Tissue Expression (GTEx) project^36^, version 8, and from BodyMap dataset^37^, respectively. We processed the data, to achieve a more comparable dataset between human and mouse, as we previously did^51^. Briefly, from the human tissues data we excluded all the pseudo-autosomal expression records and the tissues of ‘Cells EBV-transformed lymphocytes’ and ‘skin sun exposed’ and merged various brain tissues by computing the mean for each gene across them, to compare them with the available mouse brain from the BodyMap data. As for the mouse tissue data, which included 17 tissues, we merged the tissues by computing the mean for each gene in both male and female (if both existed). Using this human-mouse cross-tissue comparative data, we tested whether genes found in the *in vitro* cultured fibroblasts to be more highly expressed, are also more highly expressed in human and/or mouse tissues. To test this, we performed a one-sided Mann-Whitney test. This was done separately for human and mouse.

### Functional enrichment analysis

To study the functional enrichment of genes against the background of the entire human proteome we used g:Profiler program^24^.

### Functional analysis with the CORUM database

To study whether genes’ protein products belong to the same protein complex we used the CORUM database that is a collection of experimentally verified mammalian protein complexes^25^.

### Gene age analysis

Gene age estimations were obtained from ProteinHistorian^32^. Specifically for our analysis, we used the PPODv4-OrthoMCL and wagner1.0 parameters. For each group of genes (all DE genes in either human or mouse, and the four divergent groups of genes between human and mouse in response to dsRNA stimulation) we plotted the distribution of gene age values. A one-sided Mann–Whitney test was conducted between each one of the four groups against the DE genes to the hypothesis where the age of the DE genes is higher than the one of diverging gene groups.

### Coding sequence evolutionary analysis

Human-mouse ortholog’s percentage of identity was calculated as the mean of both the percentage of identity of the human ortholog to the mouse ortholog, and the percentage of identity of the mouse ortholog to the human ortholog. Both values were obtained from ENSEMBL^49,52^. For each group of genes (all DE genes in either human or mouse and the four divergent groups of genes between human and mouse in response to dsRNA stimulation) we plotted the distribution percentage of identity. One-sided Mann–Whitney test was conducted between each one of the four groups against the DE genes to the hypothesis where the percentage of the DE genes is higher than the one of diverging gene groups.

dN/dS ratio (non-synonymous to synonymous codon substitutions) for human and mouse orthologs was also obtained from ENSEMBL. For each group of genes (all DE genes in either human or mouse and the four divergent groups of genes between human and mouse in response to dsRNA stimulation) we plotted the distribution of dN/dS ratio. A one-sided Mann–Whitney test was conducted between each one of the four groups against the DE genes to the hypothesis where the ratio of the DE genes is lower than the one of diverging gene groups.

### Rate of gene gain and loss analysis

The significance at which a gene’s family has experienced a higher rate of gene gain and loss in the course of vertebrate evolution in comparison with other gene families (“P-value of a fast rate of gain and loss”) was obtained from ENSEMBL^52^. This statistic was calculated using the CAFE method^53^, which estimates the global birth and death rate of gene families and identifies gene families with accelerated rates of gain and loss. For each group of genes (all DE genes in either human or mouse and the four divergent groups of genes between human and mouse in response to dsRNA stimulation) we plotted the distribution of their gene gain and loss P-values. A one-sided Mann–Whitney test was conducted between each one of the four groups against the DE genes to the hypothesis where the p-value of the DE genes is higher than the one of diverging gene groups.

### Interferome analysis

From the Interferome database^38^, we downloaded the data of the differential expression analysis for relevant sets of human and mouse cell types - a total of 27 experimental sets in humans and 10 in mouse (the datasets are described below). For each human and mouse 1-to-1 ortholog, we calculated the fraction of the datasets of human, and those of mouse where the ortholog is upregulated (FC>1). We excluded from this analysis genes that were completely missing from the database (that did not appear in any of the datasets of both human and mouse).

The datasets used are as follows:

Mouse:

- one dataset of wild type murine embryo fibroblasts (MEFs) treated with 2500U IFN beta for 5h
- 5 datasets of NIH-3T3 cells treated with 100U IFN alpha, where one of them was treated for 0.5h, two replicates for 1h and the remaining two replicates for 3h
- 4 datasets of lung-whole tissue cells of BALB/c mice carrying a defective allele of the Mx1 resistance gene, treated with 10,000U of recombinant human IFN alpha A/D, one of them for 12h, one for 72h and the other two for 24h.

Human:

- two datasets of Human A549 cell line treated with IFN alpha, where one of them was treated for 6h and the other for 24h.
- one dataset of Huh7 hepatoma cells treated with IFN alpha (50 IU/ml) for 72h.
- 10 datasets of Human bronchial epithelial cells (HBECs) treated with 15-minute pulse of 1000U/ml IFN beta that each one harvested in different time points (0.25, 0.5, 1, 1.5, 2, 4, 6, 8, 12, 18 hours).
- two replicates’ datasets of Blood monocyte-derived macrophages treated with 25 ng/ml IFN alpha for 3h.
- one dataset of Human fibroblast cells isolated from umbilical veins treated with 1000U IFN alpha for 5h
- two datasets of Human endothelial cells isolated from umbilical veins, one of them treated with 1000U IFN beta for 5h, and the other with 1000U IFN alpha for 5h
- 4 datasets of Human BE(2)-C neuronal cells stimulated with IFN alpha, where two of them are replicates for 12h and the other two are replicates for 6h
- one dataset of PDOVCA#1-side population of ovarian cancer cells treated with IFN alpha2b for 5h
- one dataset of PDOVCA#1-Hoechst high cells (non-Side Population) treated with IFN alpha2b for 5h
- one dataset of monocyte derived dendritic cells treated with IFN alpha for 24h
- one dataset of monocyte derived macrophages treated with IFN alpha for 72h
- one dataset of Human hepatocyte cells treated with IFN alpha for 6h

### Promoter definition and ChIP-seq analysis of promoters of upregulated genes

Using gene annotations from ENSEMBL version 99^52^, we selected for each gene the representative transcript as the transcript with the best TSL (lowest) and if there was more than one such transcripts, we selected the longest transcript. After finding a representative transcript per gene, we defined the promoter region as the region 2,000 bp upstream of the TSS and 500 bp downstream of it, following our previous work where we investigate promoter characteristics^51^.

In the ChIP-seq analysis, we aimed to quantify the extent of agreement between the RNA and chromatin levels: between the levels of expression of genes upregulated in dsRNA-response in a species-specific manner (the four gene groups defined above), and chromatin active regions in their promoters before and after stimulation in each of the two species, as probed by ChIP-seq of H3K27ac. For example, we asked whether genes that are constitutively expressed in mouse cells, in both control and treatment and were only upregulated in response to dsRNA in human cells, will have an active H3K27 mark in both conditions in mouse but in human cells only after dsRNA-treatment.

To define promoter region as “active”, we overlapped called peaks from the ChIP-seq data (peak processing, including calling and merging, was done as described previously^11^), with promoter regions as described above, using the bedtools^54^ command: intersectBed -a {processed peaks bed file} -b {promoters bed file} -wo For each gene category (e.g., ‘mouse constitutive-high’) we computed the fraction of genes that the patterns of promoter activity agrees with gene expression across the two species in both conditions (e.g., there is a called peak in the promoters of human and mouse dsRNA condition and in the control condition of mouse, but not in the control condition of human in ‘mouse constitutive-high’ genes). To compute the enrichment of this fraction, we compared it with the fraction of all DE genes in mouse or human using Fisher’s exact test.

### Statistical analysis

Statistical analyses (Mann-Whitney test, Fisher’s exact test, Spearman’s rank-order correlation and FDR correction based on Benjamini-Hochberg procedure^55^) were performed using the SciPy package in Python (version 3.9.7). Data in boxplots represent the median, first quartile and third quartile with lines extending to the furthest value within 1.5 of the interquartile range. Plots were created using matplotlib and seaborn packages in Python, except for the PCA plots that were created using ggplot package in R.

## Declarations

### Ethics approval and consent to participate

Not applicable.

### Consent for publication

Not applicable.

### Availability of data and materials

Data used in this work is publicly available ; Arrayexpress accession - E-MTAB-5919, SRA accession: SRP120495, ArrayExpress accession: E-MTAB-5918.

## Authors’ contributions

L.S and T.H analyzed and interpreted the data. T.H wrote the manuscript with input from L.S. All authors read and approved the final manuscript.

## Acknowledgements

We would like to thank Sivan Friedman-Nakar, Irit Gat-Viks and Peter Sudmant for helpful discussions during the project and on the manuscript and Maya Lischinski for assistance with creating the initial dataset.

