## Supplemental Figures for "Evolutionary divergence of induced versus constitutive antiviral gene expression between primates and rodents"

### Supporting Information

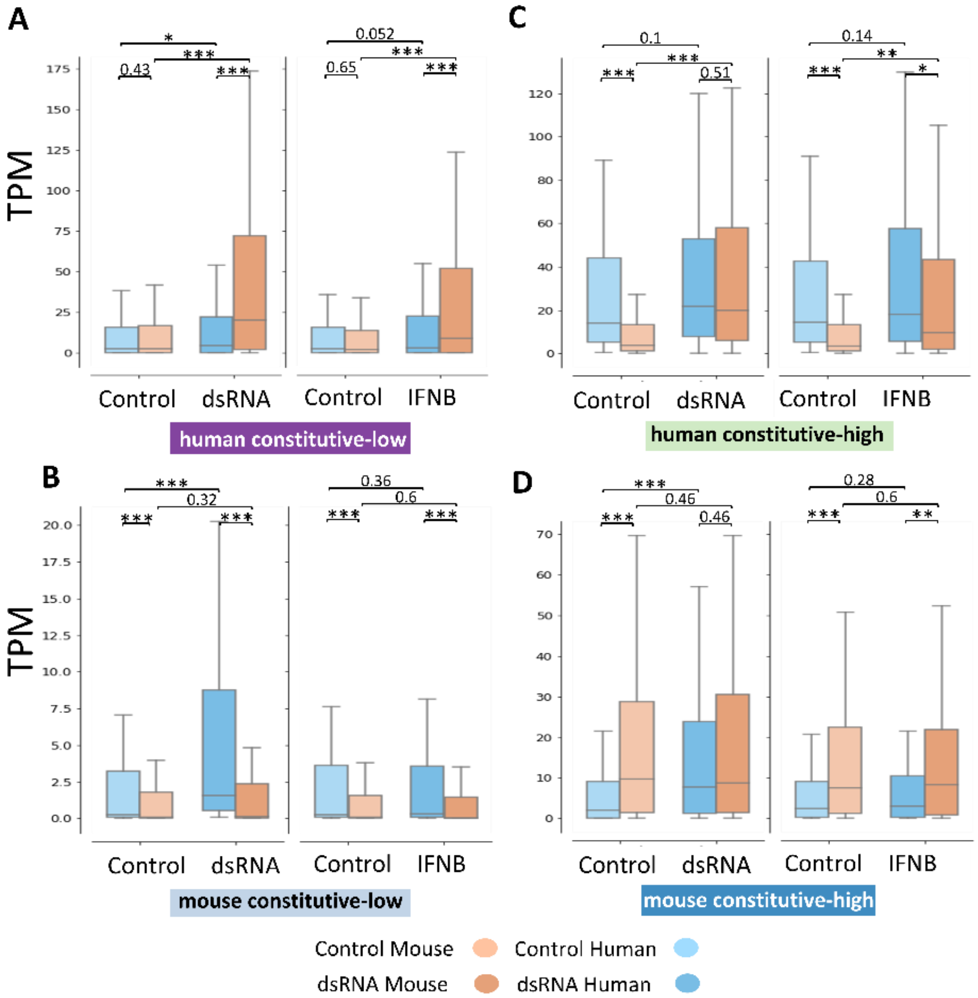


**Supporting Figure 1: Human-mouse divergence in response to dsRNA stimulation is similar to the divergence observed in response to IFN.** Averaged TPM levels of human and mouse genes in control and in stimulation conditions in both dsRNA and IFN systems for genes in (**A**) 'human constitutive-low', (**B**) mouse constitutive-low', (**C**) 'human constitutive-high' and (**D**) 'mouse constitutive-high'. FDR-corrected one- or two-sided Mann–Whitney tests were performed according to the expected pattern of expression matching to the specific group and statistical significance is shown. The observed results in A-D suggest an overall similarity in the transcriptional behavior of the human-mouse orthologs, originally identified based on their divergence in response to dsRNA, to behave similarilyy also in response to IFN.

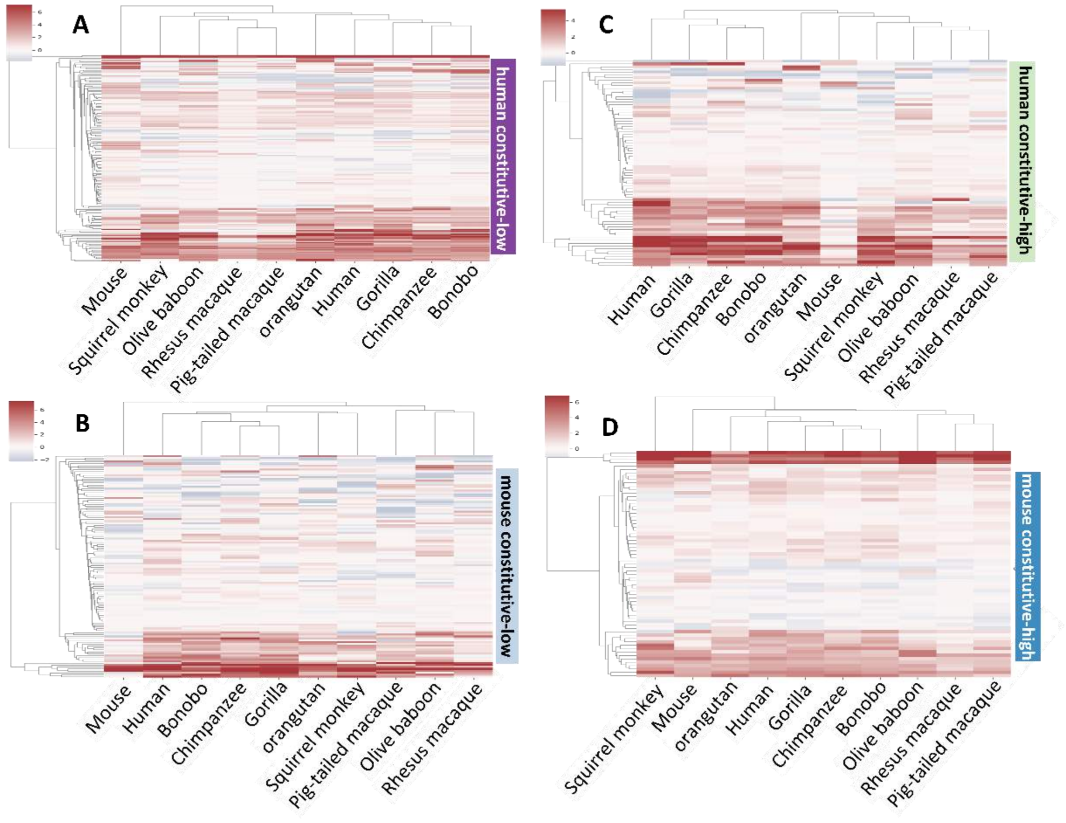


**Supporting Figure 2: Hierarchical clustering across the primate clade in the four groups of human-mouse transcriptionally divergent genes in response to dsRNA.** Hierarchically clustered heatmaps on logFC values in response to dsRNA for orthologous genes in cells from 9 primates and mouse, for each one of the 4 divergent gene groups, defined in Figure 3: (**A**) ‘human constitutive-low’, (**B**) mouse constitutive-low’, (**C**) ‘human constitutive-high’ and (**D**) ‘mouse constitutive-high’. The logFC values are from differential expression analysis between control and stimulation with dsRNA conditions for each of the species from the 10-species system (in each case, the DE values are within the same species).

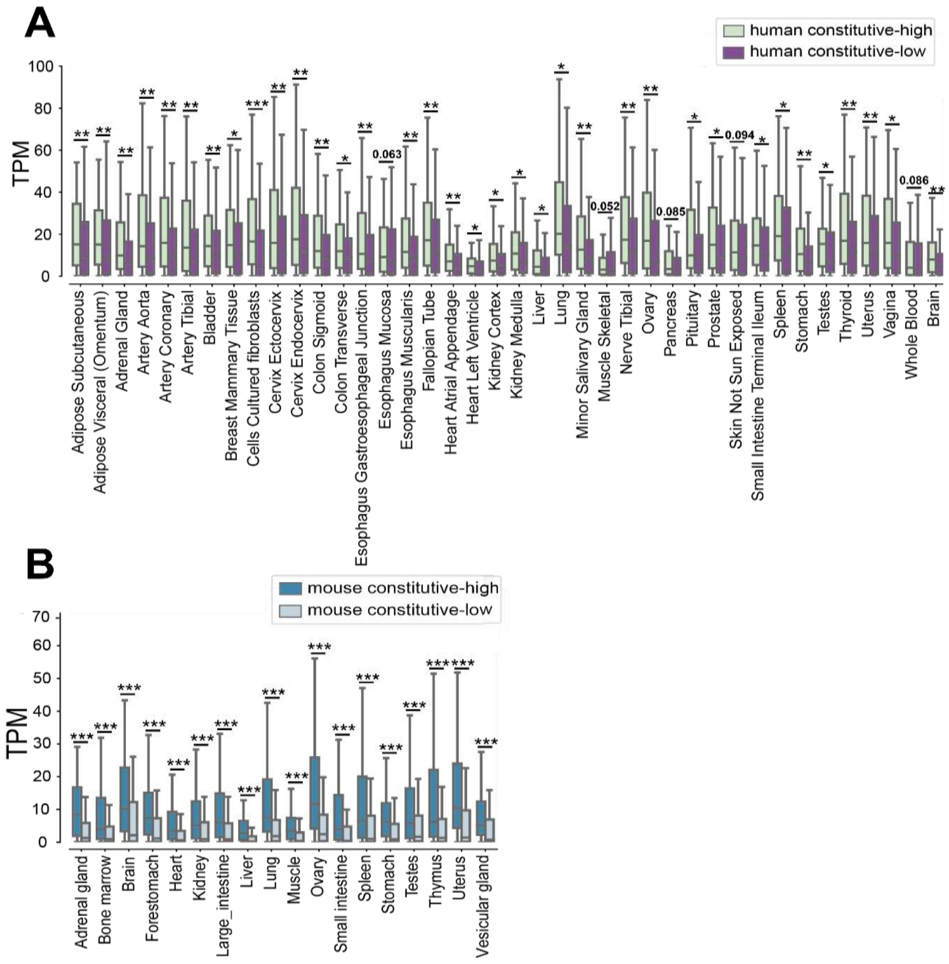


**Supporting Figure 3: Basal expression levels of constitutively low and constitutively high genes in human and mouse across tissues.** Distributions of TPM values are shown in boxplots, of constitutive-low and high genes as defined (A) for human, in 40 human tissues from the GTEx dataset[1], and (B) for mouse, in 17 mouse tissues from the BodyMap dataset[2]. FDR-corrected P-values are shown for one-sided Mann–Whitney test, performed between the basal TPM values of constitutive-high and low genes in human or mouse, for each tissue within the species. In the majority of cases (tissues / species), the constitutive-low genes are significantly lower in expression that the constitutive-high genes.

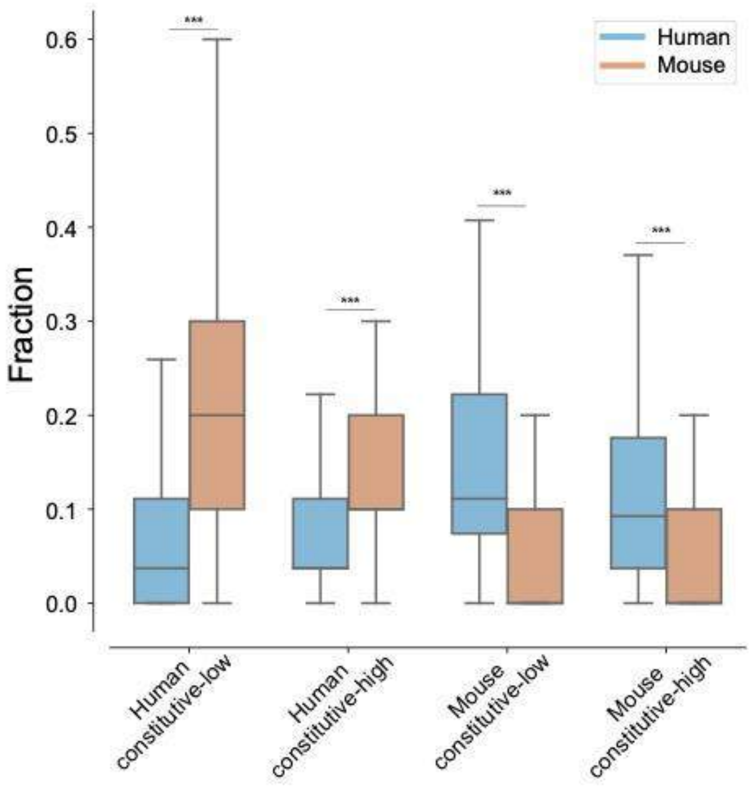


**Supporting Figure 4**: **Fraction of datasets from Interferome in which genes are upregulated in response to Interferon from the four groups of human-mouse divergent genes**. Each of the 4 divergent groups is shown, partitioned by Interferome[3] datasets of human and mouse transcriptional response data (Each dataset is based on transcriptional response to IFN in a different cell or tissue in human or mouse). FDR-corrected P-values are shown for one-sided Mann–Whitney test that was performed under the hypothesis that the fraction in human is higher (in mouse constitutive high or mouse constitutive low groups) or lower (in human constitutive high or human constitutive low groups) than in mouse. We observe that two distributions (human versus mouse) are always significantly different and that they follow our expectations based on the stimulation data from human and mouse fibroblasts. For example, in the group of genes that were identified as "human constitutive low" in human-mouse fibroblast data, we observe that the distribution of human is lower than that of mouse orthologous genes, as expected given the fact that this group is induced in mouse fibroblasts, and not in human fibroblasts, following IFN stimulation.
